# Shared and distinct neural signatures of inhibitory control deficits in attention-deficit/hyperactivity disorder and autism

**DOI:** 10.1101/2025.10.10.681732

**Authors:** Mady Roussat, Zhiyao Gao, Weidong Cai, Chandan Vaidya, Kaustubh Supekar

**Affiliations:** Department of Psychiatry & Behavioral Sciences, Stanford University, Stanford, CA USA; Wu Tsai Neurosciences Institute, Stanford University, Stanford, CA USA; Department of Psychology, Georgetown University, Washington, DC USA; Children’s Research Institute, Children’s National Medical Center, Washington, DC USA

**Keywords:** non-comorbid populations, multi-level brain analyses, neural maturation index, hyperdirect cortico-subthalamic pathway

## Abstract

**Background:** Attention-deficit/hyperactivity disorder (ADHD) and autism spectrum disorder (ASD) share deficits in inhibitory control, yet direct comparisons of their neurobiological underpinnings remain sparse and are often confounded by comorbidity. We examined shared and distinct abnormalities in brain activation and circuit connectivity associated with inhibitory control in non-comorbid ADHD and ASD.

**Methods:** Brain activity and hyperdirect pathway connectivity during inhibitory control were examined in children with non-comorbid ADHD (age: 11.1±0.24y) and children with non-comorbid ASD (age: 11.1±0.39y), compared with typically developing (TD) children (age: 10.9±0.09y), and benchmarked against healthy adults (age: 24.1±1.26y), using fMRI stop-signal task data.

**Results:** Both ADHD and ASD groups showed reduced activation in salience and frontoparietal network regions alongside increased activation in default mode network (DMN) regions, consistent with triple-network dysfunction. These activation deficits were more pronounced in ASD, whose activation profile also diverged most from adults. At the circuit level, only ASD showed abnormal hyperdirect connectivity, involving insula-STN and preSMA-STN pathways and further reflected in a composite insula/IFG-STN connectivity measure, indicating broader cortical-STN disruption. Importantly, across all children, reduced salience/frontoparietal relative to DMN engagement correlated with greater parent-reported inhibitory control deficits, and in ASD, individual differences in insula/IFG-STN connectivity showed the same association, highlighting the behavioral relevance of both network- and circuit-level abnormalities.

**Conclusions:** ADHD and ASD share network-level but differ in circuit-level abnormalities underlying inhibitory control deficits. Together, these insights advance the neurobiology of inhibitory control in ADHD and ASD and provide potential targets for neuroscience-informed interventions addressing both shared and disorder-specific mechanisms.

## Introduction

Attention-deficit/hyperactivity disorder (ADHD) and autism spectrum disorder (ASD) are common neurodevelopmental disorders that pose significant challenges in cognition and behavior. ADHD is primarily characterized by inattention, hyperactivity, and impulsivity, whereas ASD is defined by social communication difficulties and restricted, repetitive behaviors and interests (1). Despite these diagnostic distinctions, the disorders frequently overlap: approximately 20% of children with ADHD also meet criteria for ASD, and 40-80% of children with ASD exhibit ADHD symptoms (2, 3). This overlap raises the possibility that ADHD and ASD may not be entirely distinct but could represent related manifestations along a continuum. Recent research has explored this hypothesis by comparing the two disorders across various domains and revealed shared deficits in executive functions, particularly in inhibitory control, which refers to the ability to suppress prepotent or inappropriate responses in favor of goal-directed behavior (4–6). Notably, these shared deficits in inhibitory control correlate significantly with the core symptoms observed in both ADHD and ASD (7, 8). Despite the growing evidence of a common cognitive profile linking ADHD and ASD, it remains unclear whether their neurobiological underpinnings are similarly overlapping or distinct. Addressing this gap is crucial not only for understanding the pathophysiology of these disorders but also for improving diagnostic precision and developing targeted treatments that address both common and disorder-specific aspects of their neurobiology. Here we investigate these critical questions and identify both shared and distinct abnormalities in brain activation and circuit connectivity associated with inhibitory control across the two disorders. We compare children with ADHD and children with ASD against typically developing (TD) children and evaluate how these inhibitory profiles align with those observed in healthy adults.

The first aim of this study was to identify shared and distinct abnormalities in brain activation associated with inhibitory control in children with ADHD and children with ASD, using TD children as the reference group. During inhibitory control, TD children recruit a broad set of cortical and basal ganglia regions, including the anterior insula (AI), right inferior frontal gyrus (IFG), middle frontal gyrus (MFG), presupplementary motor area (preSMA), posterior parietal cortex (PPC) and the striatum (9). Investigations into neural inhibitory control deficits in children with ADHD have shown decreased activation in the insula, anterior cingulate cortex (ACC), supplementary motor areas, frontal gyrus, ventro-lateral prefrontal cortex (vlPFC), and striatum, compared with TD children (10–17). Additionally, atypical activation have been reported in the posterior cingulate cortex (PCC) and precuneus (18–20). Research on brain activation associated with inhibitory control in children with ASD, although limited, indicates that compared with TD children, children with ASD show decreased activation in the insula, PCC, precuneus and parietal regions, along with deficits in frontal, hippocampal, and occipital activation (21, 22).

While these findings suggest the potential for shared and distinct abnormalities in brain activation associated with inhibitory control in children with ADHD and children with ASD, direct quantitative comparisons between these groups remain underexplored. Here we address this critical knowledge gap using a stop signal task, which is well-suited to investigate inhibitory control (9, 23), in conjunction with functional magnetic resonance (fMRI) to systematically compare the patterns of brain activation between children with ADHD, children with ASD, and TD children. Another novel aspect of our study is that we focus on children with ADHD and children with ASD without comorbid diagnoses, thereby addressing a significant limitation of most previous studies, whose findings may have been confounded by the high rates of comorbidity between the two disorders. We hypothesized that both disorders would exhibit reduced engagement of task-positive salience network nodes, including the AI, and insufficient suppression of the task-negative default mode network nodes, including the PCC, with potential disorder-specific differences in prefrontal and basal ganglia recruitment.

The second aim of the study was to determine how the identified shared and distinct abnormalities in brain activation associated with inhibitory control in children with ASD and children with ADHD compare with those observed in healthy adults. It is well-established that during inhibitory control tasks healthy adults engage a wide range of cortical and basal ganglia nodes, including the AI, IFG, MFG, preSMA, PPC, and the striatum (9, 24). Recent findings indicate that the brain activation patterns in healthy adults during these tasks closely resemble those seen in TD children and that TD children whose brain activation patterns are more adult-like exhibit better inhibitory control abilities (9). However, it remains unclear to what extent the brain activation associated with inhibitory control in children with ADHD and children with ASD mirrors that observed in healthy adults. We addressed this question using the Neural Maturation Index (NMI), which measures how similar a child’s whole-brain activity is to that of adults. Previous studies have demonstrated the effectiveness of the NMI as a reliable marker of developmental differences in inhibitory control maturation (9). This allowed us to examine how closely brain activation patterns of inhibitory control in children with ADHD and children with ASD resemble those of healthy adults, compared with TD children and to each other. We expected both ADHD and ASD groups to have lower NMIs than TD children, indicative of less adult-like brain activation patterns and reflective of delayed developmental inhibitory control mechanisms in these clinical groups. Given the characterization of inhibitory control as a core deficit in ADHD (25), we also hypothesized that the ADHD group might show a pronounced delay characterized by a lower NMI compared with the ASD group.

The third aim of the study was to identify shared and distinct abnormalities in brain circuit connectivity associated with inhibitory control in children with ADHD and children with ASD, using TD children as the reference group. As previously noted, both TD children and healthy adults engage a similar set of cortical and basal ganglia regions during inhibitory control. The prevailing circuit-based understanding of cognitive functioning posits that these regions are key components of a cortico-basal ganglia circuit, which aids in inhibition by influencing motor control through several frontostriatal loops (26, 27) (**Figure 1f**). The direct and indirect pathways within this circuit traverse through the striatum, a critical node of the basal ganglia, modulating the thalamus to either facilitate or restrain motor actions. The hyperdirect pathway adds an additional layer to this circuitry by projecting directly from the cortex to the subthalamic nucleus (STN), bypassing the striatum and providing a rapid mechanism to modulate motor actions (27, 28). Several studies have implicated this pathway in inhibitory control (9, 23, 28, 29). While frontostriatal deficits have been characterized as part of the inhibitory control dysfunction observed in neurodevelopmental disorders (11, 12, 30), impairments in hyperdirect pathway connectivity as a potential contributor to their deficits have been overlooked in the literature. To address this gap, we examined task-modulated connectivity in this pathway in children with ADHD, children with ASD, and TD children using a seed-based generalized psychophysiological interaction (gPPI) approach. We anticipated that both the ADHD and ASD groups would exhibit reduced effective connectivity in the hyperdirect pathway compared with TD children, with potentially distinct patterns of impairments across the two disorders.

**Figure 1:**
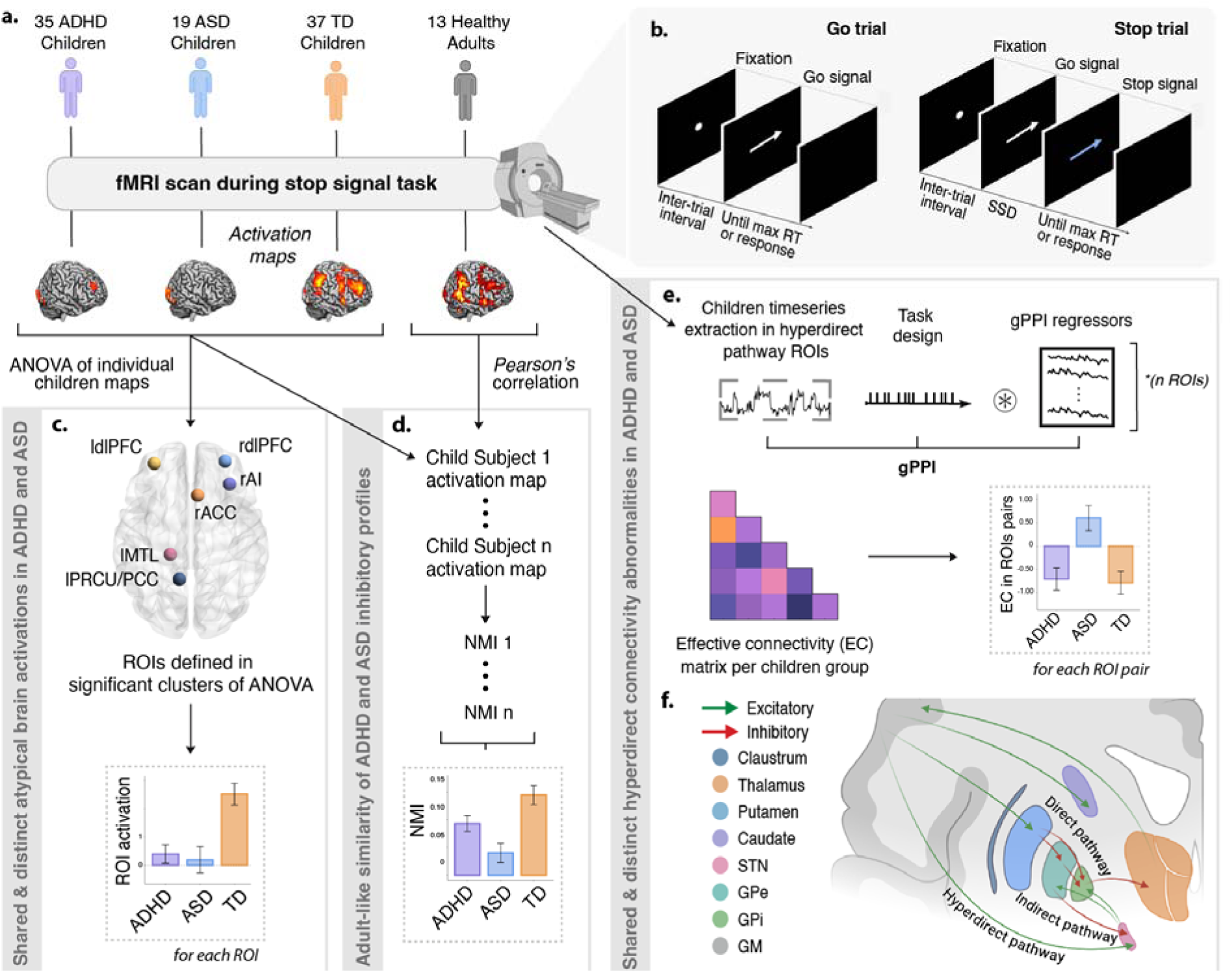
Overview of the study design and analysis pipeline. **a.** Task-based GLM analysis used to evaluate stop signal task-related brain activation in children with ADHD, children with ASD, TD children and healthy adults focusing on Successful Stop-Correct Go contrast to isolate inhibitory control. **b.** Stop signal task trial design. Each trial began with a fixation followed by an inter-trial interval. Go trials required the participant to press a button corresponding to the direction of the arrow. During Stop trials, the arrow changed color indicating that the subject should cancel their ongoing motor response. **c.** Regional level analysis of shared and distinct brain activation abnormalities during inhibitory control in children with ADHD, children with ASD and TD children, focusing on ROIs identified at peak coordinates of significant ANOVA clusters (Aim 1). **d.** Comparison of inhibitory control profiles in children with ADHD, children with ASD, and TD children, relative to healthy adults, using the neural maturation index (Aim 2). **e.** Circuit level analysis of shared and distinct effective connectivity abnormalities within the hyperdirect pathway in children with ADHD, children with ASD, and TD children using a gPPI approach (Aim 3). **f.** Schematic representation of cortico-subcortical pathways, including the hyperdirect pathway investigated in the current study. *ACC = Anterior Cingulate Cortex, ADHD = Attention-Deficit/Hyperactivity Disorder, AI = Anterior Insula, ANOVA = Analysis of Variance, ASD = Autism Spectrum Disorder, dlPFC = Dorso-lateral Prefrontal Cortex, EC = Effective Connectivity, fMRI = Functional Magnetic Resonance Imaging, GLM = General Linear Model, GM = Grey matter, GPe = external globus pallidus, GPi = internal globus pallidus, gPPI = Generalized Psychophysiological Interaction, IFG = Inferior Frontal Gyrus, MFG = Middle Frontal Gyrus, MTL = Medial Temporal Lobe, NMI = Neural Maturation Index, PCC = Posterior Cingulate Cortex, PRCU = Precuneus, PreSMA = Pre-supplementary Motor Area, ROI = Region of Interest, RT = Reaction Time, SS = Successful Stop, SSD = Stop signal delay, STN = Subthalamic Nucleus, TD = Typically Developing*

## Results

### Participant characteristics

The child groups did not differ significantly in age (F(2,41.111)=0.270, p=.765, η^2^= 0.008) or in mean/maximum scan-to-scan displacement (F(2,88)=1.146, p=.323, η^2^=0.025 and F(2,60.889)=1.168, p=.316, η^2^=0.027, respectively) (**Table 1**). Descriptive statistics for adults were excluded from comparisons as their data were used solely for the NMI reference map.

**Table 1:** Descriptive statistics of ADHD, ASD and healthy participants. Age and head motion parameters were compared between child groups using one-way analysis of variance (ANOVA) and a two-sided chi-squared test was used to compare their sex distributions. Age and head motion parameters are reported as mean ± standard error of the mean. χ *test statistic for chi-square test, F test statistic for one-way ANOVA; ADHD = Attention-Deficit/Hyperactivity Disorder, ASD = Autism Spectrum Disorder, df = Degrees of Freedom, NS = Not Significant, p = P-value, TD = Typically Developing*.

|  | Healthy Adults<br>(n = 13) | ADHD Children<br>(n = 35) | ASD Children<br>(n = 19) | TD Children<br>(n = 37) | Between group comparison<br>(children groups) |  |  | Post-hoc tests |  |
| --- | --- | --- | --- | --- | --- | --- | --- | --- | --- |
| | | | | | $\chi^2$ | $df$ | $p$ | Groups | $p$ |
| Sex (male/female) | 10/3 | 22/13 | 19 males | 22/15 | 10.77 | 2 | 0.005 | TD - ASD; ADHD - ASD | 0.001; 0.002 |
| | | | | | $F$ | $df$ | $p$ | Groups | $p$ |
| Age (in years) | 24.1±1.26 | 11.1±0.2 | 10.9±0.4 | 10.9±0.1 | 0.27 | 2 | 0.765 | NS | NS |
| **Head motion parameters (mm)** |  |  |  |  |  |  |  |  |  |
| *Range* |  |  |  |  |  |  |  |  |  |
| Mean scan-to-scan displacement | 0.07±0.01 | 0.14±0.02 | 0.13±0.03 | 0.11±0.01 | 1.146 | 2 | 0.323 | NS | NS |
| Max scan-to-scan displacement | 0.44±0.08 | 1.49±0.21 | 1.42±0.26 | 1.12±0.12 | 1.168 | 2 | 0.316 | NS | NS |

### Differences in parent-reported executive functioning, including inhibitory control, among children with ADHD, children with ASD, and TD children

Parent-reported executive functioning, as measured by the Behavior Rating Inventory of Executive Function (BRIEF), differed significantly across groups. A one-way ANOVA revealed significant main effects of group on all BRIEF domains (all p’s < .05; **Table 2**). Post hoc comparisons showed that both children with ADHD and children with ASD had largely higher normalized t-scores than TD children. Importantly, significant elevations were observed in the inhibitory control domain, consistent with its central role in both disorders. No significant differences emerged between ADHD and ASD groups, indicating that while the underlying neural mechanisms may diverge, children with ADHD and ASD share comparably elevated deficits in inhibitory control, relative to TD children.

**Table 2:** Neuropsychological BRIEF assessments summary of children with ADHD, children with ASD and TD children. Each subscale of BRIEF was compared between child groups using one-way analysis of variance (ANOVA) and normalized t-scores are reported as mean ± standard error of the mean. *F test statistic for one-way ANOVA; ADHD = Attention-Deficit/Hyperactivity Disorder, ASD = Autism Spectrum Disorder, BRIEF =* Behavior Rating Inventory of Executive Function, *df = Degrees of Freedom, p = P-value, TD = Typically Developing*.

|  | ADHD Children<br>(n = 35) | ASD Children<br>(n = 19) | TD Children<br>(n = 37) | Between group comparison<br>(children groups) |  |  | Post-hoc tests |  |
| --- | --- | --- | --- | --- | --- | --- | --- | --- |
|  |  |  |  | *F* | *df* | *p* | Groups | *p* |
| BRIEF (Normalized T-Scores) |  |  |  |  |  |  |  |  |
| Shift | 65.2±2.3 | 70.3±2.8 | 53.3±2.2 | 11.297 | 2 | <0.001 | ADHD > TD; ASD > TD | <0.001; 0.001 |
| Inhibitory Control | 63.1±1.8 | 63.4±2.9 | 52.4±1.7 | 9.549 | 2 | <0.001 | ADHD > TD; ASD > TD | 0.002; <0.001 |
| Emotional Control | 61.9±1.9 | 61.5±2.9 | 54.1±2.3 | 3.848 | 2 | 0.026 | ADHD > TD | 0.031 |
| Initiate | 66.3±1.8 | 65.2±2.5 | 52.6±2.2 | 13.174 | 2 | <0.001 | ADHD > TD; ASD > TD | <0.001; <0.001 |
| Working Memory | 70.5±1.5 | 66.1±2.9 | 51.3±2.9 | 20.007 | 2 | <0.001 | ADHD > TD; ASD > TD | 0.002; <0.001 |
| Plan/Organize | 69.0±1.4 | 65.5±2.5 | 51.4±2.9 | 17.605 | 2 | <0.001 | ADHD > TD; ASD > TD | 0.002; <0.001 |
| Organization of Materials | 65.1±1.7 | 60.6±2.3 | 49.2±2.1 | 18.742 | 2 | <0.001 | ADHD > TD; ASD > TD | 0.001; <0.001 |
| Monitor | 65.4±1.3 | 62.0±2.4 | 50.2±2.2 | 19.599 | 2 | <0.001 | ADHD > TD; ASD > TD | 0.001; <0.001 |

### Differences in brain activation during inhibitory control among children with ADHD, children with ASD, and TD children

The within-group activation map contrasting Successful Stop and Correct Go conditions showed significant activation clusters in the ADHD group, including the right anterior insula (rAI), right dorsolateral prefrontal cortex (rdlPFC), bilateral occipital poles, and the fusiform gyrus. Children with ASD showed significant activation in the left AI, bilateral occipital poles, and the fusiform gyrus. The TD group showed widespread significant activation clusters, including the right anterior cingulate cortex (ACC), right caudate, bilateral frontal and temporal gyri, bilateral PPC, bilateral AI, and bilateral frontal and occipital poles (**Figure 2**).

**Figure 2:**
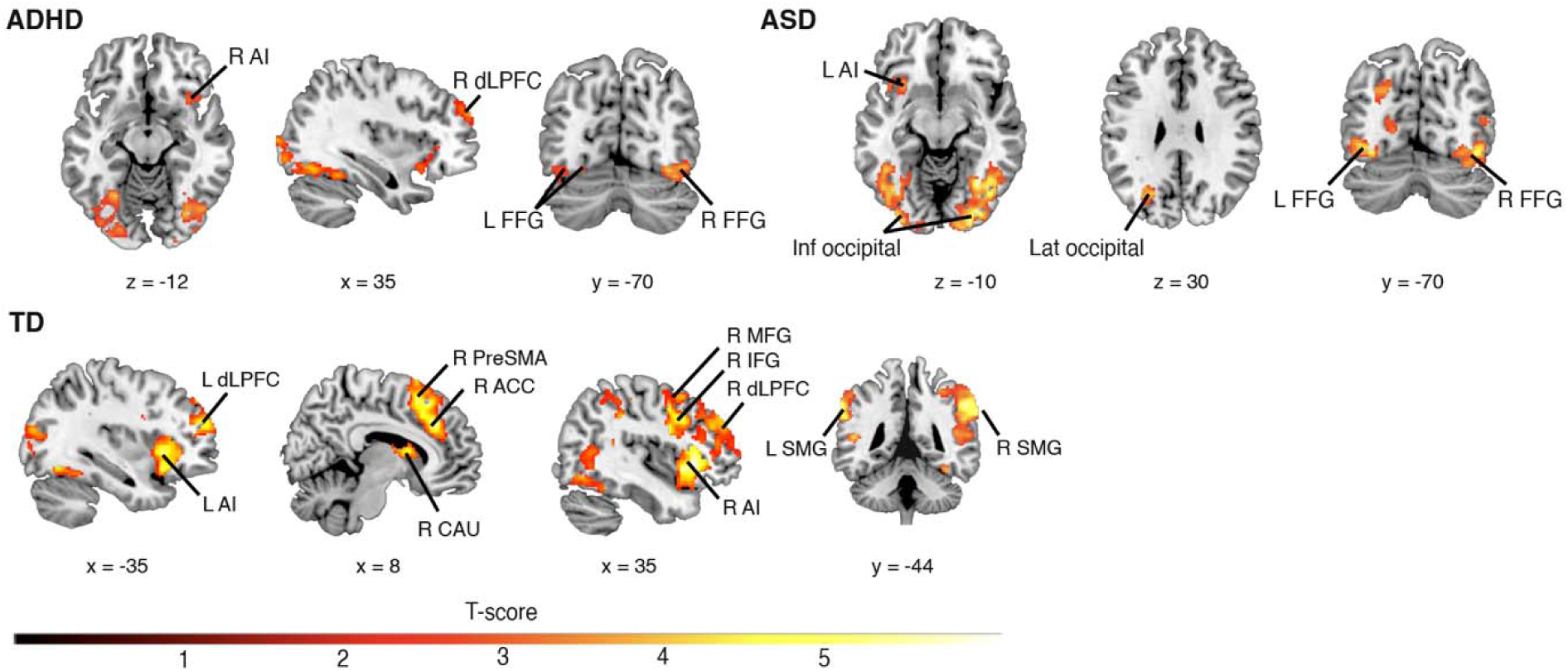
Brain activation during inhibitory control in children with ADHD, children with ASD, and TD children. Within group activation map contrasting Successful Stop and Correct Go conditions in ADHD, ASD, and TD child groups (height threshold p < 0.01, cluster extent p < 0.05, spatial extent = 100 voxels)

A one-way between-groups ANOVA revealed significant main effects of group in six clusters centered on the right AI, right ACC, left Precuneus/PCC, left medial temporal lobe (MTL), right dlPFC, and left dlPFC (**Figure 3a**; **Table 3**). ROI analyses showed that TD children had significantly greater activation during inhibitory control in SN nodes including right AI and right ACC than children with ADHD (*rAI:* p<.001, *rACC:* p=.047) and children with ASD (*rAI*: p=.001, *rACC:* p<.001) (**Figure 3b**, **Table 4**). Conversely, both ADHD and ASD groups exhibited significantly greater activation in the left Precuneus/PCC (p=.003 and p=.041, respectively) and MTL (both p’s<.001) than TD children, suggesting reduced engagement of the task-positive salience network and inadequate suppression of the task-negative DMN in both disorders. Additionally, TD children showed significantly higher frontal activation than children with ADHD and children with ASD including left dlPFC (p=.045 and p<.001, respectively) and right dlPFC (p=.011 and p<.001, respectively). The ADHD group also showed significantly higher activation than the ASD group in both right ACC (p=.015) and right dlPFC (p=.039).

**Figure 3:**
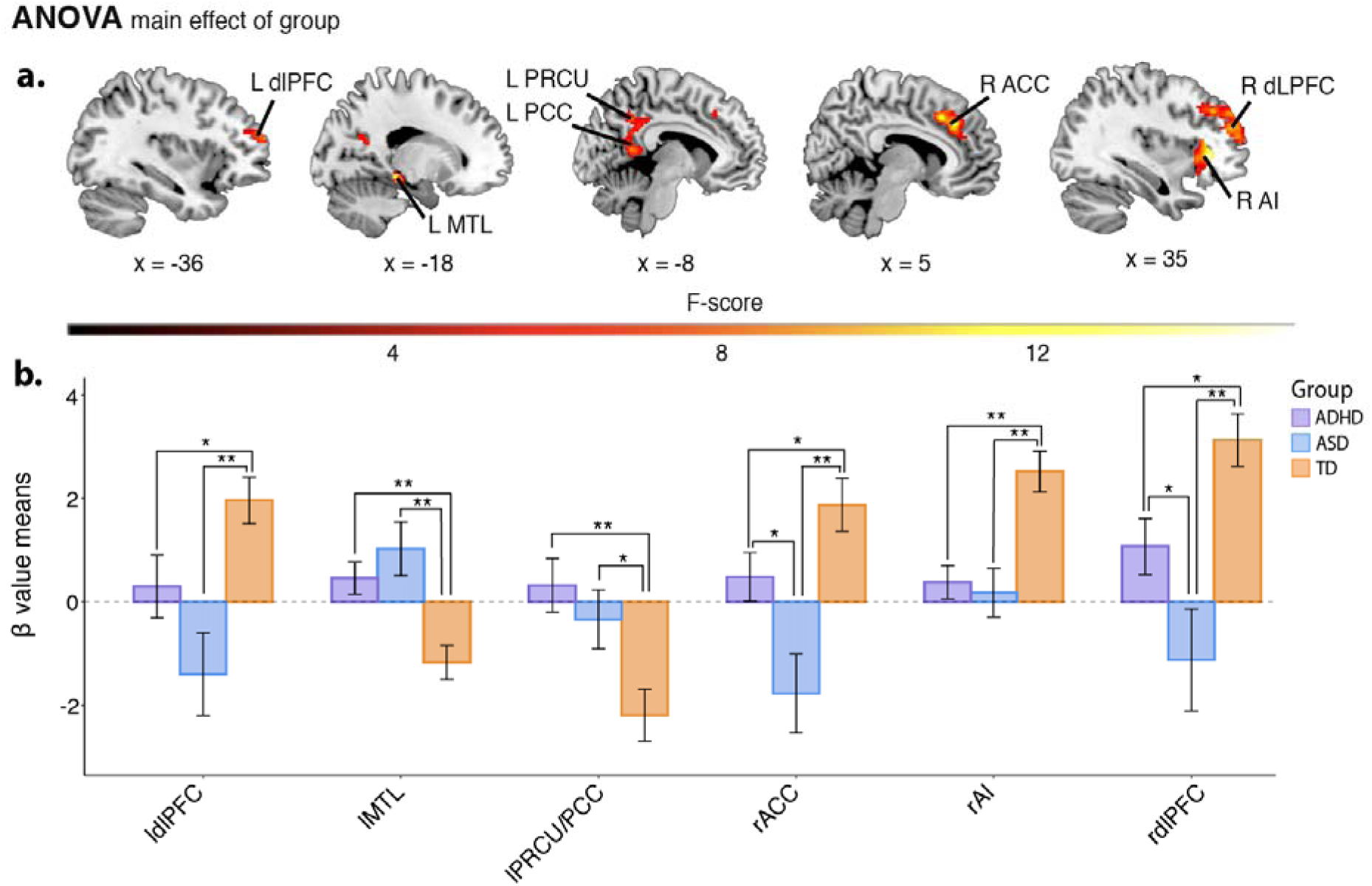
Differences in brain activation during inhibitory control among children with ADHD, children with ASD and TD children. a. One-way ANOVA revealed regional activation differences across six clusters within the salience, fronto-parietal and DMN networks between ADHD, ASD, and TD groups during inhibitory control (height threshold p < 0.01, cluster extent p < 0.05, spatial extent = 100 voxels). b. Comparative bar plots of activation level (represented by β values means) in ROIs identified at peak coordinates of significant ANOVA clusters between ADHD, ASD and TD groups. *\*\* p < .01, * p < .05*

**Table 3:**
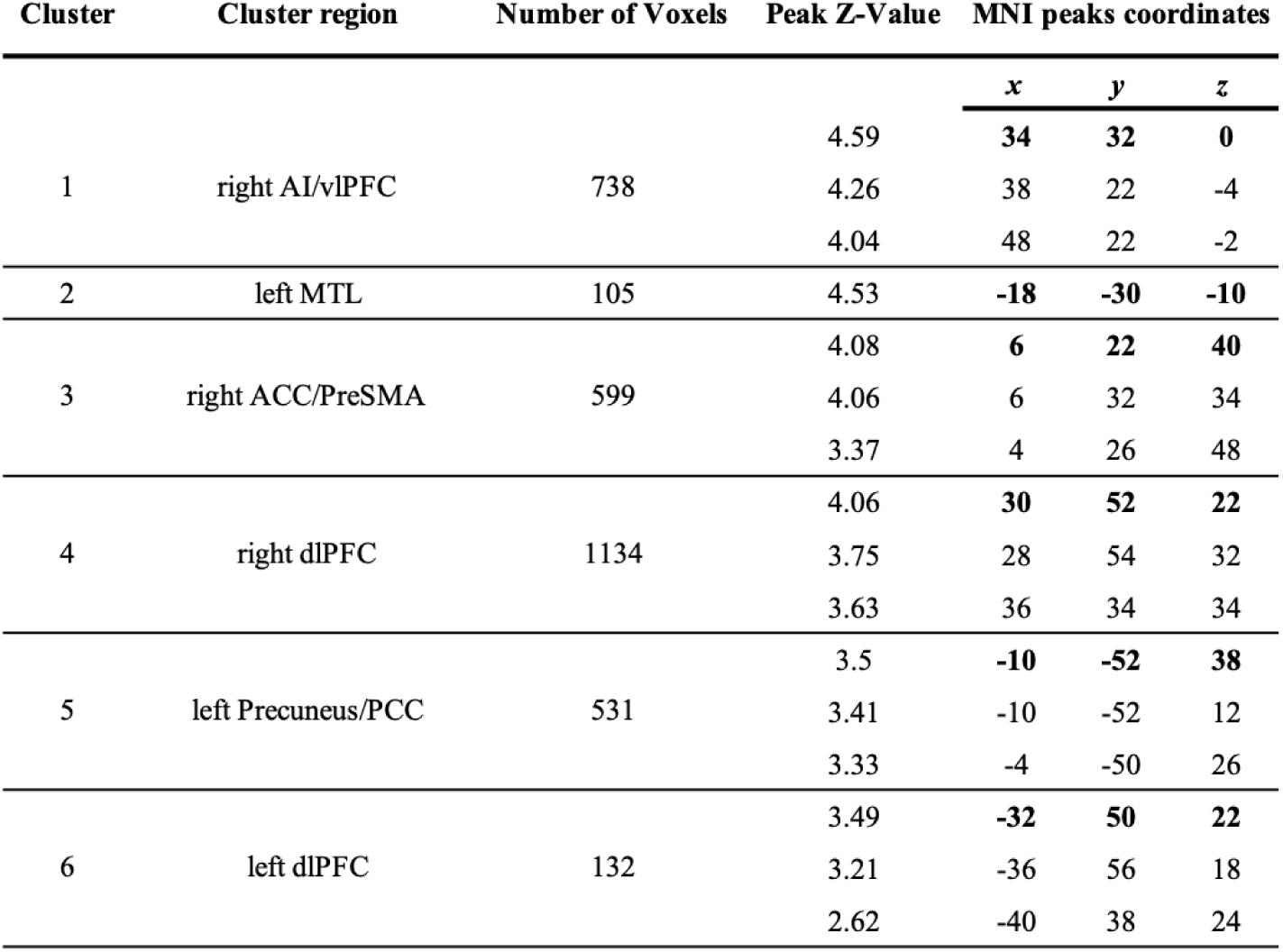
Summary table of significant brain activation revealed by between-group ANOVA comparison (children with ADHD, children with ASD and TD children) contrasting Successful Stop versus Correct Go trials. (height threshold p < 0.01, cluster extent p < 0.05, spatial extent = 100 contiguous voxels). *ACC = Anterior Cingulate Cortex, AI = Anterior Insula, dlPFC = Dorso-lateral Prefrontal Cortex, MTL = Medial Temporal Lobe, PCC = Posterior Cingulate Cortex, PreSMA = Pre-supplementary Motor Area, vlPFC = Ventro-lateral Prefrontal Cortex*.

| Cluster | Cluster region | Number of Voxels | Peak Z-Value | MNI peaks coordinates |  |  |
| --- | --- | --- | --- | --- | --- | --- |
|  |  |  |  | *x* | *y* | *z* |
| 1 | right AI/vlPFC | 738 | 4.59 | **34** | **32** | **0** |
|  |  |  | 4.26 | 38 | 22 | -4 |
|  |  |  | 4.04 | 48 | 22 | -2 |
| 2 | left MTL | 105 | 4.53 | **-18** | **-30** | **-10** |
| 3 | right ACC/PreSMA | 599 | 4.08 | **6** | **22** | **40** |
|  |  |  | 4.06 | 6 | 32 | 34 |
|  |  |  | 3.37 | 4 | 26 | 48 |
| 4 | right dlPFC | 1134 | 4.06 | **30** | **52** | **22** |
|  |  |  | 3.75 | 28 | 54 | 32 |
|  |  |  | 3.63 | 36 | 34 | 34 |
| 5 | left Precuneus/PCC | 531 | 3.5 | **-10** | **-52** | **38** |
|  |  |  | 3.41 | -10 | -52 | 12 |
|  |  |  | 3.33 | -4 | -50 | 26 |
| 6 | left dlPFC | 132 | 3.49 | **-32** | **50** | **22** |
|  |  |  | 3.21 | -36 | 56 | 18 |
|  |  |  | 2.62 | -40 | 38 | 24 |

**Table 4:**
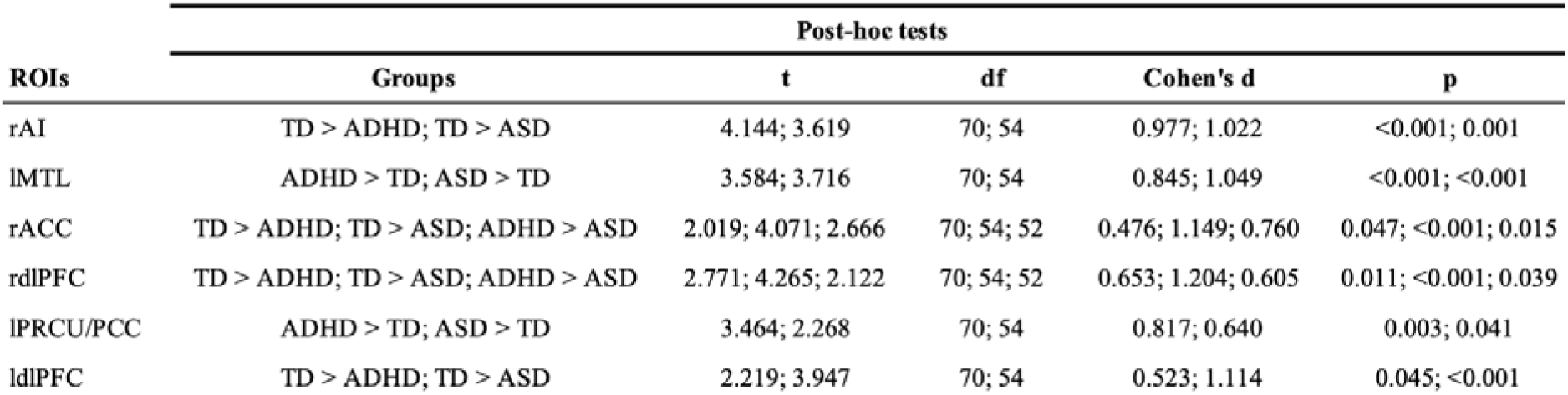
Statistical summary table of ROI-level analysis of brain activation contrasting Successful Stop versus Correct Go trials between ADHD, ASD, and TD groups. *t test statistic for post-hoc tests; ACC = Anterior Cingulate Cortex, ADHD = Attention-Deficit/Hyperactivity Disorder, AI = Anterior Insula, ASD = Autism Spectrum Disorder, df = Degrees of Freedom, dlPFC = Dorso-lateral Prefrontal Cortex, MTL = Medial Temporal Lobe, p = P-value, PCC = Posterior Cingulate Cortex, PRCU = Precuneus, ROI = Region of Interest, TD = Typically Developing, ** p < .01, * p < .05*

### Comparison of inhibitory control activation profiles in children with ADHD, children with ASD, and TD children relative to healthy adults

We next investigated how the identified shared and distinct abnormalities in brain activation associated with inhibitory control in children with ASD and children with ADHD compared with those observed in healthy adults. We addressed this question using the NMI, which measures how similar a child’s whole-brain activity is to that of adults. This analysis revealed significant group differences (F(2, 88)=8.134, p<.001, η^2^=0.156; **Figure 4**). TD children had the highest NMI, indicating more adult-like activation patterns, significantly different from both the ADHD group (p=.02, t(70)=2.226, d=0.525) and the ASD group (p=.001, t(54)=3.797, d=1.072). Moreover, the ASD group had a significantly lower NMI than the ADHD group (p=.029, t(52)=-2.244, d=-0.639), suggesting a more pronounced deviation from healthy adult brain activation patterns in the ASD group.

**Figure 4:**
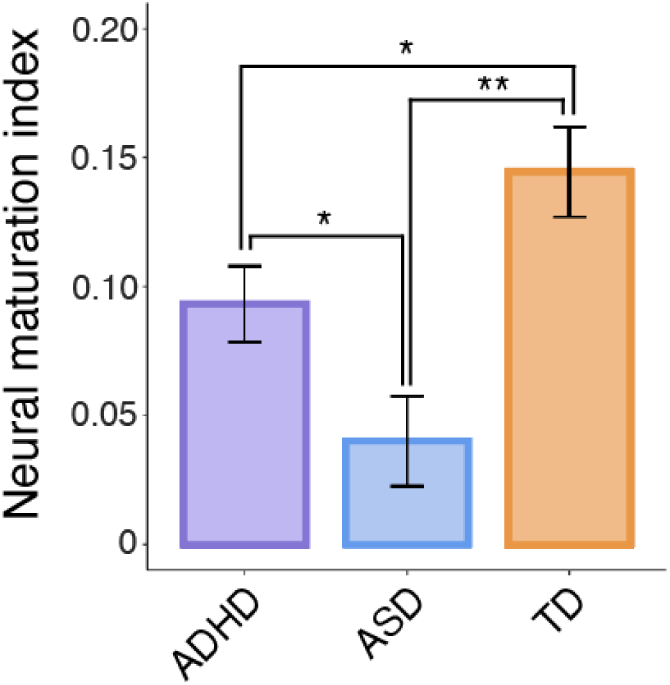
Differences in the neural maturation index among children with ADHD, children with ASD and TD children. ADHD = Attention-Deficit/Hyperactivity Disorder, ASD = Autism Spectrum Disorder, TD = Typically Developing, ** p < .01, * p < .05

### Differences in effective connectivity of the hyperdirect pathway during inhibitory control among children with ADHD, children with ASD, and TD children

We next examined brain circuit connectivity associated with inhibitory control in children with ADHD and children with ASD, using TD children as the reference group. Specifically, we examined connectivity of the hyperdirect pathway, a key brain circuit critical for the execution of inhibitory control, using a gPPI-based effective connectivity approach. This analysis revealed significant group differences in effective connectivity during inhibitory control, specifically in hyperdirect links rAI-lSTN (F(2, 85.072)=5.604, p=.028, η^2^=0.098) and rPreSMA-lSTN (F(2, 88)=7.047, p=.012, η^2^=0.138) (**Figure 5**). Post-hoc pairwise comparisons showed significant differences between ASD and both TD and ADHD groups in rAI-lSTN effective connectivity (p=.0095, t(54)=2.840, d=0.871 and p=.0092, t(52)=3.099, d=0.927, respectively) and rPreSMA-lSTN (p=.002, t(54)=3.561, d=1.041 and p=.002, t(52)=3.421, d=1.002, respectively), whereas no significant differences were found between TD and ADHD groups. Extending beyond these individual connections, we computed a composite insula/IFG-STN index to capture convergent cortical input to the STN from key salience and inhibitory control nodes. This measure also differed across groups (F(2,85.072)=4.001, p=.02, η²=0.1), with ASD again significantly different from both TD (p=.023, t(54)=2.28, d=0.68) and ADHD (p=.002, t(52)=3.20, d=0.91).

**Figure 5:**
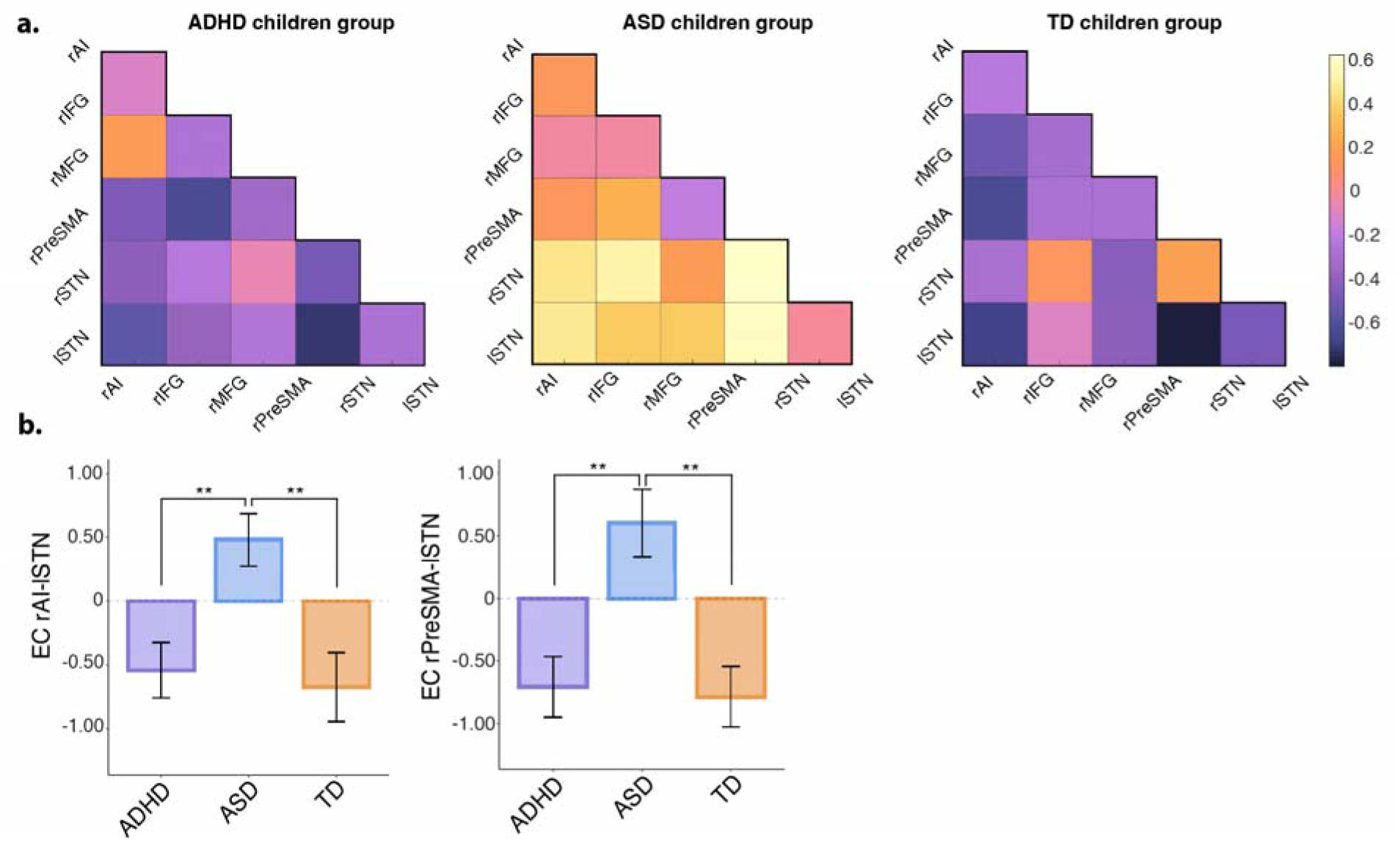
Differences in the effective connectivity within the hyperdirect pathway among children with ADHD, children with ASD and TD children. **a.** Effective connectivity matrix per child group **b.** Comparative bar plots of rAI-lSTN and rPreSMA-lSTN effective connectivity between ADHD, ASD and TD groups during inhibitory control. *\*\* p < .01, * p < .05*

### Brain-behavior relationships

To establish behavioral relevance, we examined associations between neural measures and parent-reported inhibitory difficulties. At the network level, we computed a balance index reflecting task-positive versus task-negative engagement, defined as the difference between average beta values in salience/frontoparietal and DMN regions (for the Successful Stop versus Correct Go contrast), given the central role of triple-network coordination in inhibitory control. Across all children, reduced salience/frontoparietal relative to DMN engagement was significantly associated with greater inhibitory deficits on the BRIEF Inhibitory control scale (Spearman’s ρ=-0.34, p=.002). At the circuit level, after statistically controlling for variance explained by the network-level balance index, composite insula/IFG-STN connectivity remained significantly associated with inhibitory deficits on the BRIEF Inhibitory control scale in the ASD group (Spearman’s ρ=-0.50, p=.02); no such association was observed in ADHD or TD groups.

### Control analyses

We conducted two separate control analyses to ensure that our results were not confounded by head motion. First, we applied our analysis pipeline to a subset of children with ADHD and children with ASD using more stringent movement exclusion criteria. Second, we conducted ANCOVA instead of ANOVA, using mean scan-to-scan displacement as a covariate of no interest to account for potential motion-related confounds. In both cases, all key findings replicated.

## Discussion

To our knowledge, this study is among the first to examine shared and distinct abnormalities in both brain activation and circuit connectivity associated with inhibitory control in non-comorbid children with ADHD and ASD. Across analyses, we identified convergent disruptions in large-scale cognitive control networks in both groups as well as disorder-specific deficits most pronounced in ASD. Specifically, relative to TD peers, both ADHD and ASD children showed abnormal engagement of salience, frontoparietal, and default mode networks during stopping, accompanied by reduced similarity to adult activation profiles. Crucially, only ASD children showed additional deficits in salience and frontoparietal nodes as well as impaired hyperdirect cortico-subthalamic connectivity, suggesting more pronounced or qualitatively different impairments in the circuitry that implements inhibitory control in ASD.

### Shared and distinct abnormalities in brain activation associated with inhibitory control in children with ADHD and children with ASD

#### Shared abnormalities

We identified shared abnormalities in brain activation associated with inhibitory control in children with ADHD and children with ASD, compared to TD children, within the triple-network model of cognitive control. This model posits that the salience network detects salient events and orchestrates cognitive control by engaging the frontoparietal network and suppressing the default mode network in response to cognitively demanding tasks (7, 31). Consistent with this model, TD children showed robust salience and frontoparietal networks engagement alongside DMN deactivation during stopping. In contrast, both children with ADHD and children with ASD exhibited a breakdown in triple-network coordination: reduced activation in salience and frontoparietal network regions (AI, ACC, and dlPFC) coupled with greater activation in DMN regions (precuneus/PCC and MTL). Abnormalities also extended to canonical inhibitory control regions (23, 24), including the vlPFC, which is consistently implicated in ADHD and increasingly in ASD (32, 30), as well as the preSMA, which supports motor response inhibition. Importantly, across all children, reduced salience/frontoparietal engagement relative to DMN engagement correlated with greater parent-reported inhibitory deficits, highlighting the behavioral relevance of the observed shared network-level dysfunction.

#### Distinct abnormalities

Beyond these shared disruptions, we observed distinct abnormalities in brain activation associated with inhibitory control in children with ASD, who exhibited significantly reduced activation in the ACC node of the salience network and the dlPFC node of the frontoparietal network compared to both children with ADHD and TD children. This contrasts with two prior studies that reported increased activation in these same regions in ASD relative to ADHD (30, 33). This discrepancy may stem from methodological differences, including the inclusion of comorbid cases and wider developmental ranges in prior work, whereas our study focused on non-comorbid children. Notably, our observation of decreased activation in the ACC and dlPFC in ASD aligns with meta-analytic evidence (32, 34). ADHD children, by contrast, did not show these additional cortical deficits, and we did not detect basal ganglia abnormalities frequently reported in prior studies (11, 12, 30), possibly due to small effect sizes or the limited sensitivity of univariate approaches. Taken together, these results reinforce emerging evidence that triple-network dysfunction represents a transdiagnostic signature of neurodevelopmental disorders (35), while also pinpointing specific regional vulnerabilities in salience and frontoparietal networks that distinguish ASD from ADHD.

### Developmental profile of brain activation associated with inhibitory control in children with ADHD and children with ASD

Our next aim was to determine how the identified shared and distinct abnormalities in brain activation associated with inhibitory control in children with ADHD and children with ASD compare with activation patterns observed in healthy adults. To do so, we used the NMI, a validated metric that assesses the maturity of inhibition-related brain mechanisms by comparing children’s brain activation patterns to those of adults performing the same inhibitory task (9). As expected, TD children showed the most adult-like activation profiles, whereas both ADHD and ASD groups diverged significantly. Importantly, children with ASD had significantly lower NMI than children with ADHD, indicating that their activation pattern was farthest from the adult reference.

These findings suggest that ADHD and ASD may follow different developmental trajectories. ADHD has often been characterized as a delay in the maturation of cognitive control networks, with children with ADHD eventually reaching typical development milestones but on a slower timetable (36–39). In this framework, the ADHD brain activation pattern during inhibitory control may reflect a maturational lag rather than a qualitative deviation. In contrast, ASD has been associated with divergent developmental trajectories, reflected in phenomena such as early brain overgrowth, atypical synaptic pruning, and anomalous cortical trajectories (40, 41). Our observation that ASD activation patterns were least adult-like supports this deviance perspective, indicating that inhibitory control networks in ASD may not simply be delayed but may follow a qualitatively different course. Because our data are cross-sectional and the NMI provides an indirect measure of developmental maturity, longitudinal studies are needed to confirm whether the observed patterns reflect delays or qualitative deviations in developmental trajectories.

### Shared and distinct abnormalities in brain circuit connectivity associated with inhibitory control in children with ADHD and children with ASD

Our final aim was to identify shared and distinct abnormalities in the hyperdirect pathway, a rapid cortical-subthalamic circuit critical for inhibitory control (9), in children with ADHD and children with ASD.

#### Shared abnormalities

No shared abnormalities in hyperdirect connectivity during inhibitory control were observed between ADHD and ASD, suggesting divergent circuit-level mechanisms of inhibitory control despite overlapping network dysfunction.

#### Distinct abnormalities

Children with ASD showed distinct abnormalities in hyperdirect connectivity along two links, AI-STN and preSMA-STN, during inhibitory control compared to both ADHD and TD groups. The anterior insula is central for detecting salient cues and initiating network switching (24, 31), while the preSMA contributes to adaptive response selection and inhibitory engagement via its rapid connections to the STN (42–44). Disruption of both AI-STN and preSMA-STN pathways suggests that in ASD the relay of stop commands from salience detection and motor control regions to the basal ganglia is compromised, potentially weakening the efficiency of rapid inhibitory responses.

Extending beyond these individual links, a composite insula/IFG-STN index, capturing convergent cortical input to the STN from both salience and canonical inhibitory control hubs, was also significantly different in ASD. The IFG is a core inhibitory region, and when combined with the insula, this measure reflects the integration of salience detection and inhibitory implementation signals into the basal ganglia (45, 46). At the behavioral level, composite insula/IFG-STN connectivity correlated with inhibitory deficits in ASD, even after accounting for variance explained by the network-level balance index. This indicates that circuit-level abnormalities in ASD provide unique explanatory power beyond brain network-level dysfunction. Together, these findings offer circuit-level evidence that inhibitory dysfunction in ASD reflects not only impaired signaling from single cortical regions but also disrupted coordination of convergent cortical inputs to the STN. In contrast, children with ADHD showed preserved hyperdirect connectivity, which did not differ from TD children. This suggests that inhibitory failures in ADHD may arise not from breakdown of the stopping mechanism itself but from inconsistent engagement of otherwise intact circuits, consistent with accounts linking ADHD to variability in attention and network recruitment rather than to impaired motor-stopping circuitry.

Overall, our hyperdirect connectivity findings of potential underutilization of intact hyperdirect circuits in ADHD and disruption of cortical-STN communication in ASD illustrate a circuit-level dissociation between ADHD and ASD, suggesting that disorders with overlapping inhibitory control deficits may be underpinned by distinct neural circuit mechanisms.

#### Clinical implications

The shared and distinct neural abnormalities in inhibitory control identified here suggest potential avenues for translational relevance. Shared dysfunction across salience, frontoparietal, and default mode networks points to transdiagnostic targets, suggesting that interventions aimed at strengthening task-positive control and suppressing task-negative activity may benefit both ADHD and ASD. Concurrently, disorder-specific patterns highlight the need for tailored approaches. In ADHD, preserved hyperdirect connectivity indicates intact stopping circuitry but inconsistent engagement, suggesting interventions, such as stimulant medication or cognitive training, that enhance salience detection and network recruitment may remediate inhibitory deficits. In ASD, hypoactivation of salience and frontoparietal regions combined with disrupted AI-STN and preSMA-STN connectivity indicates atypical circuitry that may require compensatory strategies, such as neuromodulation or targeted training to strengthen cortical-basal ganglia communication. More broadly, recognizing both the shared network-level dysfunction and the disorder-specific circuit vulnerabilities is critical for advancing precision psychiatry approaches to inhibitory control deficits in ADHD and ASD.

## Conclusions

This study provides a comprehensive characterization of both shared and disorder-specific neural mechanisms of inhibitory control in ADHD and ASD. We found that children with these neurodevelopmental disorders exhibit overlapping disruptions in key large-scale brain networks, marked by reduced recruitment of salience and frontoparietal network regions alongside insufficient suppression of the default mode network. These convergent abnormalities suggest shared network-level dysfunctions that may underlie executive function impairments observed across both disorders. The neural profiles of ADHD and ASD were, however, not identical. Children with ASD showed more pronounced abnormalities and specific deficits in the hyperdirect cortico-subthalamic inhibitory pathway, which were not evident in ADHD. Crucially, these network- and circuit-level abnormalities were behaviorally relevant, linking directly to parent-reported inhibitory difficulties. Collectively, these findings refine our neurobiological understanding of inhibitory control in ADHD and ASD and highlight the need for targeted, disorder-specific interventions that address both shared network dysfunctions and distinct circuit-level abnormalities. Moreover, these findings provide a foundation for future investigations of comorbid presentations, which are associated with greater symptom severity, poorer treatment outcomes, and reduced adaptive functioning compared with either disorder alone (35).

## Methods and Materials

### Study Design

Participants from all cohorts underwent an fMRI scan during which they performed a behavioral stop signal task. The Behavior Rating Inventory of Executive Function (47) (BRIEF) assessment was completed by legal guardians for children but was not collected for adults. Informed consent was obtained from participants or legal guardians, and studies were approved by the IRBs of the respective institutions.

### Participants

#### Georgetown cohort

322 children were recruited from Children’s National Hospital (CNHS) clinics at Georgetown University. Participants were 8-14 years old, had a full-scale IQ>70, and no other medical conditions or contraindications to fMRI. Diagnoses of ADHD and ASD were based on the DSM-IV-RT / DSM-5 (1, 48) and confirmed using clinical assessments: the Mini-International Neuropsychiatric Interview for ADHD (49) and the Autism Diagnostic Observation Schedule-Generic for ASD (50). Stimulant treatments were withheld for at least 24 hours prior to scanning. 100 children had a complete dataset for analysis (imaging and behavioral data). After preprocessing, 9 children were excluded due to excessive motion (described below) and 1 child due to unusable behavioral data. Of the remaining subjects, 76 had a primary diagnosis of ADHD or ASD and 22 were excluded due to comorbid diagnoses of ADHD and ASD. Therefore, the study included 35 subjects with non-comorbid ADHD (age: 11.1±0.24y, 22M/13F) and 19 subjects with non-comorbid ASD (age: 11.1±0.39y, 19M).

#### Stanford cohort

78 children were recruited through local schools at Stanford University. They had no history of psychiatric or neurological disorders and were not taking stimulants. After preprocessing, 33 children were excluded due to excessive motion (described below), 1 for missing data, 5 for poor stop signal task performance and 2 due to incomplete scans. The study therefore included 37 TD children (age: 10.9±0.09y, 22M/15F).

#### OpenfMRI cohort

fMRI data with the same stop signal task design as Georgetown and Stanford data were obtained from 13 healthy adults (age: 24.1±1.26y, 10M/3F) from the public OpenfMRI database (https://openneuro.org/datasets/ds000008). Detailed information about the data is available in its reference paper (29).

### Stop signal task

Children completed one (children with ADHD or ASD) or two runs (TD children) of 96 trials of a stop signal task (**Figure 1b**). During Go trials, participants responded to an arrow by pressing a left or right button corresponding to its direction. During Stop trials, the arrow changed color after a time delay indicating that subjects should cancel their ongoing response. Participants were presented with a majority of Go trials, creating a prepotency for Go responses and effectively measuring their ability to inhibit a prepotent response on Stop trials. Trials began with a fixation followed by a jittered inter-trial interval. The stop signal delay (SSD) between Go and Stop signal was adjusted based on performance: +50ms after Successful Stop, -50ms after Unsuccessful Stop.

#### Georgetown cohort

Imaging data were acquired on a Siemens Trio 3T scanner, except for 15 participants scanned on a Siemens Prisma_fit 3T with identical parameters after a scanner update. Functional images of 37 axial slices (thickness ∼3mm) were acquired in an ascending and interleaved order using a T2*-weighted gradient-echo pulse sequence with the parameters: repetition time (TR) = 2000ms, echo time (TE) = 30ms, flip angle = 90°, matrix size = 64x64, field of view (FOV) = 256x256mm and voxel size = 3x3x3mm^3^. 145 volumes were obtained for each participant over the full task run (∼4-5 minutes). Structural images of 176 slices (thickness 1mm) were acquired using a high-resolution T-1 weighted magnetization-prepared rapid acquisition gradient-echo with the parameters: TR = 1900ms, TE = 2.52ms, flip angle = 90°, matrix size = 256x256, FOV = 256x256mm and voxel size = 1x1x1mm.

#### Stanford cohort

Imaging data were collected on a 3T GE Signa scanner using an 8-channel head coil. Functional images of 29 axial slices (thickness ∼4mm) were acquired in an ascending and interleaved order using a T2*-weighted gradient-echo spiral in-out pulse sequence with the parameters: TR = 2000ms, TE = 30ms, flip angle = 80°, matrix size = 64x64, FOV = 220x220mm and voxel size = 3.4x3.4x4mm^3^. 174 volumes were obtained for each participant over a full task run (∼5-6 minutes). Structural images of 166 slices (thickness 1mm) were acquired using a high-resolution T-1 weighted spoiled-gradient-recalled inversion recovery three-dimensional MRI sequence with the parameters: TR = 8.4ms, TE = 1.8ms, flip angle = 15°, matrix size = 256x192, FOV = 22cm and voxel size = 1x1x1mm.

#### OpenfMRI cohort

Functional images of 33 slices (thickness ∼4mm) were acquired on a 3T Siemens AG Allegra scanner in an ascending and interleaved order using a T2*-weighted echo-planar sequence with the following parameters: TR = 2000ms, TE = 30ms, flip angle = 90°, matrix size = 64x64, FOV = 200x200mm and voxel size = 3x3x4mm. 166 volumes were acquired for each participant over a full task run (∼5-6 minutes).

### Imaging data analysis

*For all data analysis, assumptions of normality and homogeneity of variance were checked prior to conducting analysis of variance (ANOVA). When the homogeneity assumption was violated, a Brown-Forsythe correction was applied. Multiple comparisons were corrected by controlling the False Discovery Rate (FDR) with the Benjamini-Hochberg procedure*.

Preprocessing with SPM12 included quality control, slice-timing correction, realignment, normalization to MNI space and 6mm Gaussian smoothing, plus co-registration with skull-stripped anatomical scans for children. Healthy children and adults were excluded if their mean frame displacement (FD) exceeded 0.5mm and/or their max FD exceeded 5mm in any run. Due to lower fMRI compliance in clinical groups (51), slightly less stringent motion criteria were applied for Georgetown subjects, excluding those with movement parameters over 10 mm and/or mean FD over 0.5 mm.

#### Task-based GLM analysis

A general linear model (GLM) design was used to evaluate task-related brain activation in children with ADHD, ASD, TD children and healthy adults. Four conditions Correct Go, Incorrect Go, Successful Stop, and Unsuccessful Stop were modeled as events using the onsets of the Go stimuli, convolved with the hemodynamic response function (HRF) and entered as regressors in the GLM. Temporal derivatives of the HRF were included in the model and 6 movement parameters were added as covariates of no interest. The contrast Successful Stop-Correct Go was generated to isolate inhibitory control and used to create a t-statistics map for each individual and group activation maps (**Figure 1a)**.

#### Regional activation differences analysis

To examine shared and distinct abnormalities in brain activation associated with inhibitory control in children with ADHD, children with ASD and TD children (reference group), their first-level GLM contrast images (contrast Successful Stop-Correct Go) were entered together into a one-way ANOVA to estimate regional activation differences between groups during inhibitory control (**Figure 1c**). The resulting map was thresholded (p<.01, spatial extent=100, Monte Carlo simulation approach), and regions of interest (ROIs) were defined as 6mm spheres using MarsBaR at peak coordinates of significant ANOVA clusters. GLM-derived β-values were extracted from each ROI for each subject (contrast Successful Stop-Correct Go) and compared between child groups using post hoc t-tests.

#### Neural maturation index analysis

We used the NMI to assess how the inhibitory profiles of children with ADHD, ASD, and TD children identified in the first aim of our study compared to those observed in healthy adults. This index measures the developmental maturation of inhibition-related brain mechanisms by comparing a child’s whole-brain activity to that of adults during inhibitory control (9). Adults first-level contrast images were entered into a group-level analysis to create a reference map of inhibitory control activation, which was thresholded (p < .01, spatial extent = 100), and significant voxels were included in a mask. For each child, we extracted voxel t-values from their individual activation map (contrast Successful Stop-Correct Go) using the created mask, calculated the Pearson correlation of the t-values with adult reference values, and applied a Fisher transformation to obtain the NMI. NMIs were compared between child groups using a one-way ANOVA and post hoc t-tests (**Figure 1d)**.

#### Effective connectivity analysis

Seed-based gPPI (52) was used to assess effective connectivity differences in the hyperdirect pathway between ADHD, ASD and TD children. Seeds were defined as 6mm spheres (4mm for STN) centered on the rAI, rIFG, rMFG, right pre-supplementary motor area (rPreSMA), rSTN, and lSTN. This independent set of ROIs was selected from prior meta-analytic work (9) to reflect canonical adult inhibitory control regions, allowing us to investigate hyperdirect connectivity differences in children from a developmental perspective. The gPPI design included psychological factors representing the main effect of task (onset times of Correct Go, Incorrect Go, Successful Stop, and Unsuccessful Stop conditions convolved with the HRF), a physiological factor (time course of a seed) and the PPI interaction regressors (element-by-element product of the psychological and physiological factors). Inhibition-related modulation effect on effective connectivity was estimated for each individual by subtracting the PPI regressor β-values between the Successful Stop and Correct Go conditions. Values were averaged by seed pairs (e.g. rAI-lSTN and lSTN-rAI) and compared between groups using one-way ANOVAs and post hoc t-tests (**Figure 1e**).

#### Brain behavior relationships analysis

To assess the behavioral relevance of neural abnormalities, we examined associations between brain measures of inhibitory control and parent-reported executive functioning. Parent ratings were obtained using the BRIEF, focusing on the Inhibitory control subscale as a validated index of everyday inhibitory difficulties. At the network level, we derived a balance index reflecting task-positive versus task-negative engagement, computed as the difference in average GLM beta values (Successful Stop - Correct Go contrast) between salience/frontoparietal and DMN regions identified in the group-level ANOVA. At the circuit level, we computed a composite insula/IFG-STN connectivity index by averaging effective connectivity estimates from rAI-lSTN and rIFG-lSTN hyperdirect pathways, capturing convergent cortical input to the subthalamic nucleus from salience and inhibitory control regions. To isolate circuit-specific contributions, we performed partial correlation analyses in which variance explained by the network-level balance index was covaried out. All correlations between neural indices and BRIEF Inhibitory control scores were tested using Spearman’s rank correlation coefficients to reduce sensitivity to outliers. Group-specific correlations were examined separately in children with ADHD, children with ASD, and TD children, as well as across all children combined.

## Financial Disclosures

All authors report no biomedical financial interests or potential conflicts of interest.

## Acknowledgments

This research was supported by a grant from the National Institutes of Health (LM014597), a NARSAD Young Investigator Award, and a Stanford Innovator Award, to K.S, and by a grant from the National Institutes of Health (MH124816) to W.C.

## Notes

### Competing Interest Statement

The authors have declared no competing interest.

